# The rapid degradation of translated upstream regions points to an inefficient translation initiation process

**DOI:** 10.1101/2024.11.25.625198

**Authors:** Michael L. Tress

## Abstract

Large-scale experimental analyses find ever more abundant evidence of translation from start codons upstream of the canonical start site. This translation either generates entirely new proteins (from novel upstream open reading frames) or produces isoforms with extended N-terminals when the novel start codon is in frame

Most extended N-terminals are likely to just add a disordered region to the canonical protein isoform, but some may also block the recognition of the signal peptide causing the isoform to accumulate in the incorrect cellular compartment. This analysis finds evidence that upstream translations that would interfere with signal peptides are detected in expected quantities in ribosome profiling experiments, but that the equivalent N-terminally extended protein isoforms are significantly reduced in multiple proteomics experiments.

This suggests that these isoforms are likely to be degraded shortly after translation by the ubiquitination pathway, thus preventing the build up of potentially harmful proteins with hydrophobic regions in the cytoplasm. In addition, this is further evidence that most of the transcripts translated from upstream start sites are the result of an inefficient translation initiation process. This has implications for the annotation of proteins given the huge numbers of upstream translations that are being detected in large-scale experiments.

## Introduction

Translation from regions upstream of canonical start codons has been shown to be commonplace [1-4], and in particular in-frame upstream translations that produce isoforms with extended N-terminals have abundant proteomics support [4,6]. Evidence suggests that the upstream translation detected in these experiments is only the tip of the iceberg [4]. This translation has two main characteristics that distinguish it from translation from canonical coding exons. Firstly, most start codons are non-canonical [2-5], and secondly, with a few notable exceptions [4], these upstream regions have little detectable conservation signal [3,4].

Translation from upstream of the canonical ATG can be divided into three main types [4]. Open reading frames (ORFs) that have both their initiation codon and stop codon upstream of the canonical ATG (uORFs), ORFs that upstream of principal ATG and overlap coding exons in a different frame (overlapping uORFs, or uoORFs), and 5’ extensions that initiate before the ATG and read through to the coding exons in the same frame. Translated uORFs and uoORFs produce proteins that are very different from the canonical proteins, and 5’ extensions generate proteins that are identical to the canonical protein but with a longer N-terminal. There is considerably more protein evidence for translation from 5’ extensions than for other types of upstream translation [5,6].

The GC content of the 5’ UTR that include translated upstream regions is remarkably high [4], so in the case of the N-terminal extensions, the extra section of protein sequence at the N-terminal is likely to be disordered and in most cases will not have much effect on the function of the canonical isoform. That means that even if an N-terminal extension is produced as a result of an inefficient translation process, the isoform may not be harmful to the cell, especially if it is translated in relatively small quantities, as appears to be the case [4]. However, there is at least one situation in which an N-terminal extension has the potential to alter protein function drastically: if the N-terminal extension blocks a signal peptide.

Signal peptides are N-terminal hydrophobic sequences that are required for translocation across the endoplasmic reticulum membrane [7]. Localization of the secreted and trans-membrane proteins that pass through the endoplasmic reticulum is under the control of signal recognition particles. These bind to ribosomes from the start of the translation process, and if the nascent protein emerging from the ribosome is recognised as a signal peptide, translation is halted and the signal recognition particle delivers the nascent protein along with ribosome to the Sec61 complex [8,9]. From where the protein is translated directly into the endoplasmic reticulum lumen [8,9]. However, if the protein does not have a signal peptide at the N-terminal, the signal recognition particle detaches from the ribosome [10]. A mostly disordered N-terminal extension of a protein that precedes a signal sequence will probably cause the signal recognition particle to detach early, leaving the translated protein in the cytoplasm, the wrong cellular compartment [11].

Proteins must be in the appropriate cellular compartment. If they accumulate, mislocalized proteins will affect cell function and homeostasis by interacting with proteins vital to the organelle, and interactions between mislocalised and cytosolic proteins can eventually lead to aggregation and even lead to neurodegeneration [12,13]. In the cytoplasm, the exposed hydrophobic regions of mislocalised trans-membrane and secretory proteins are recognised by the *BAG6* complex [11, 14], leading to their polyubiquitination and degradation [15].

Here, multiple proteomics experiments provide clear evidence that protein isoforms with elongated N-terminals ahead of signal peptides are likely to be mislocalized and to be degraded via the ubiquitination pathway [11].

## Methods

### Signal peptide prediction

Signal peptides were predicted for coding genes as part of the APPRIS database [16] based on scores from SignalP (v4.1, 17). The APPRIS signal peptide module generates a score between -4 and 4 for each protein isoform. The APPRIS database assumes that any protein with a score of 2 or more has a signal peptide. However, for this analysis isoforms with an APPRIS signal peptide of score of 1 were also included because peptides with this score also agreed with the predicted or experimental subcellular location annotated by UniProtKB [18]. Scores of 0 or lower had much less agreement with UniProtKB sub-cellular location annotations. Signal peptides were predicted for the principal isoforms of all GENCODE v36 coding genes [19]. APPRIS principal isoforms were used as the reference because they have been shown to be the best approximation of the canonical protein isoform [20, 21].

The same analysis of signal peptides was also carried out using the SignalP predictions from the Human Protein Atlas [22]. The Human Protein Atlas predicts signal peptides using SignalP 6.0 [17] which predicts more types of signal peptide. However, the Human Protein Atlas does not include all the GENCODE v36 coding genes. The results were almost identical to the APPRIS analysis.

### Fedorova *et al*. analysis

Upstream translations were taken from the supplementary materials of this analysis [3]. All upstream translations were included in the study, excluding those that were not protein coding in GENCODE v36. They were separated into those that had protein evidence and those that only had ribosome profiling support based on the data from the supplementary materials.

### Zhu *et al*. analysis

Upstream translations were again listed in the supplementary materials of this analysis [2]. The results from both the A431 cell lines and normal tissues were analysed. All upstream translations (tagged by the authors as “N-terminal extensions”) were included in the study. Those that were not protein coding in GENCODE v36 were excluded. All N-terminal extensions were supported by peptides.

### Rodriguez *et al*. analysis

Upstream translations were listed in the supplementary materials of this analysis [4] and tagged as “N-terminal extension” by the authors. All upstream translations were protein coding in GENCODE v36. All N-terminal extensions were supported by peptides.

### PeptideAtlas analysis

Genes with novel upstream translations that had peptide evidence from PeptideAtlas [23] in this analysis [6] were provided by the investigators.

### Chen *et al*. analysis

Upstream 5’ extensions were listed in the supplementary materials in this analysis [24] and tagged as “extension”. Those that were not protein coding in GENCODE v36 were excluded. All extensions were supported by ribosome profiling data.

### Annotated N-terminal extensions

The set of genes that had alternative isoforms with N-terminal extensions that were not conserved across primates were provided by the investigators of the Rodriguez *et al*. paper [4].

## Results

### The C1Q-like family

Two of the large-scale experiments in this analysis [3, 4] confirm the presence of translated upstream regions in members of the C1Q-like (C1QL) family. Transcript evidence for upstream translation in *C1QL2* and *C1QL3* was reported in one of the ribosome profiling experiments [3], and an N-terminal extension in *C1QL4* had proteomics support [4].

C1Q-like (C1QL) proteins are secreted proteins found across all vertebrates. *C1QL1, C1QL2* and *C1QL3* are mostly expressed in nervous tissues and *C1QL1* and *C1QL3* mediate cell adhesion through the *ADGRB3* receptor [25, 26]. In adult mice trans-synaptic interaction between *C1ql1* and *Adgrb3* has been shown to promote the elongation of climbing fibers in Purkinje cells [27, 28]. Less is known about *C1QL2* and *C1QL4*, except that *C1QL4* is testis-expressed rather than brain-expressed. Jawed vertebrate species have all four C1QL proteins.

C1Q-like (C1QL) genes clearly undergo upstream translation. In *C1QL4*, the upstream region, translated from a highly conserved ATT codon, is clearly under purifying selection across mammals [4]. The ATT codon is conserved in the other three ancient C1QL genes and in the ancestor of the C1QL family in lamprey [Figure 1], and the upstream regions of each C1QL gene have conserved basic residues and a conserved valine and alanine-rich region immediately prior to the canonical methionine [Figure 1].

**Figure 1.**
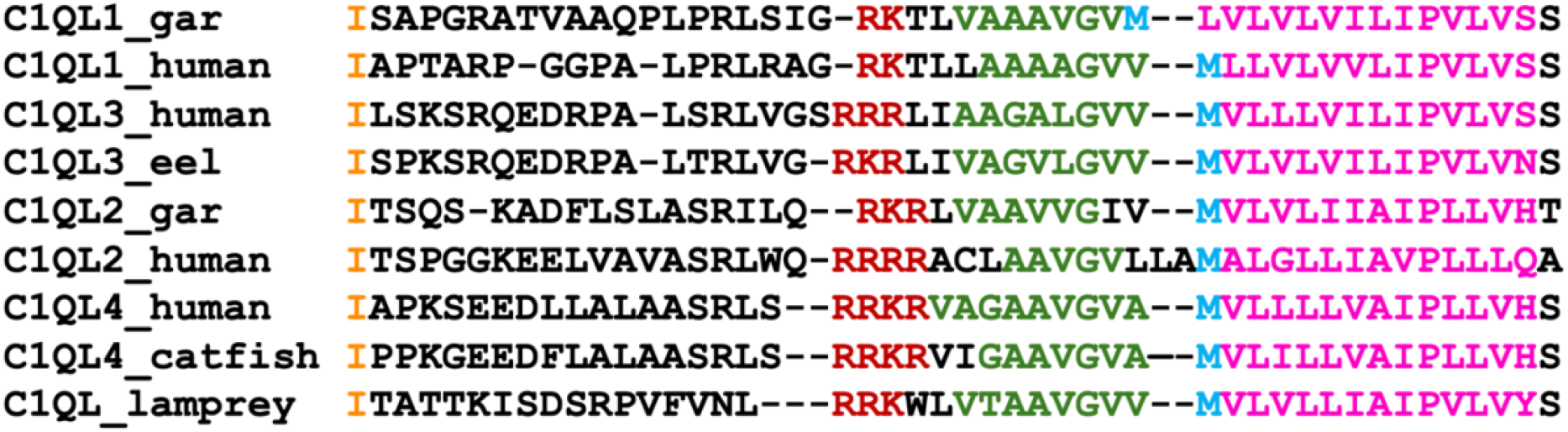
Alignment of C1Q-like proteins from human, fish and lamprey. The position of the conserved ATT start codon is marked as an isoleucine in orange (even though it probably codes for a methionine), a conserved basic motif in dark red, and a conserved alanine, valine and glycine-rich region in green. The canonical ATG is shown in blue and the hydrophobic region of the predicted signal peptide in pink.

Canonical C1QL proteins are supposed to be secreted and have signal peptides. It is known that at least *C1QL1* and *C1QL3* function as secreted proteins. However, signal peptides must start at the N-terminal end of the protein sequence [10], so C1QL isoforms with extended N-terminals will not be secreted because the signal peptide will not be recognised. Although proteins that build up in the incorrect cellular compartment are expected to interfere with the cellular processes, the ancient origins and clear cross-species conservation of the C1QL extended N-terminals shows that two populations of C1QL proteins (secreted and non-secreted) are meant to exist. The conserved VGA region in C1QL extensions is predicted to extend the signal peptide helix [29], and all have a conserved basic region next to the predicted extended helix.

### Does the loss of signal peptides lead to degradation?

The translation of the upstream region of *C1QL4* is supported by protein evidence. Since the extended isoform cannot be secreted, it must have a different role from the canonical isoform. One question is whether or not this example can be extrapolated to all signal peptide blocking N-terminal extensions. If the *C1QL4* example is typical, we would expect to detect peptides for other upstream regions, showing that other extended N-terminals that precede signal peptides also accumulate in the cytoplasm. However, if there is little or no peptide evidence for these regions, it strongly suggests that these N-terminal extensions are degraded to protect the cell because they could interfere with cellular processes.

Signal peptides were mapped using the annotations from the APPRIS database [see methods]. Over the whole GENCODE v36 reference set (with read-through genes eliminated [30]), 14.7% of the genes had signal peptides [Figure 2], while the percentage (13.8%) was slightly lower for those 14,888 genes that were detected across the five large-scale proteomics experiments [4].

**Figure 2.**
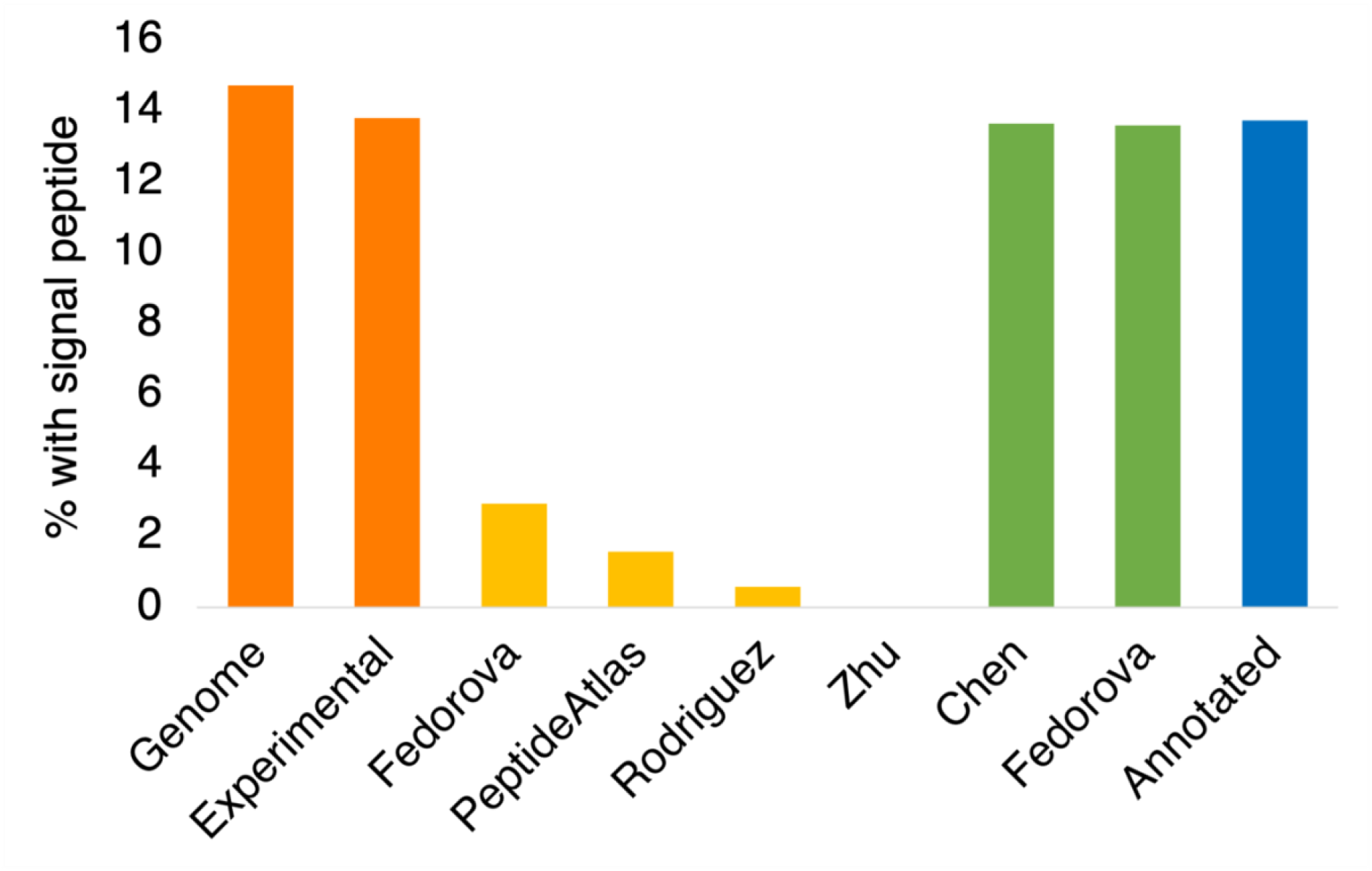
Percentage of genes with signal peptides predicted via APPRIS. The percentage of different sets of genes that have signal peptides in their principal isoform [16] according to the SignalP [17] predictor in APPRIS [16]. In orange, the percentage for the whole gene set (Genome) and for those genes detected by 5 large-scale proteomics experiments in Rodriguez et al (Experimental). In yellow, the percentage of genes with peptides for their translated upstream regions that had signal peptides in the Fedorova [3], PeptideAtlas [6], Rodriguez [4] and Zhu [2] analyses. In green, the percentage of genes with ribosome profiling evidence for upstream translation that have signal peptides in the Fedorova [3] and Chen [24] analyses. In blue, the percentage of genes with non-conserved annotated upstream translations (Annotated, [4]) that have signal peptides.

The Rodriguez *et a*l. analysis [4] detected peptides for N-terminal extensions in 170 genes. If N-terminal extensions are not regulated, 23 of these genes (13.8%) would be expected to have signal peptides in their principal isoforms. However, the only gene predicted to have a signal peptide in its principal isoform that had supporting peptides for its upstream region was *C1QL4*. Just one in 170 genes (0.59%) is much lower than expected and a Chi-squared test indicated that this was highly significant (p < 0.005).

The *C1QL4* isoform is highly conserved and is almost certainly not in a different cellular compartment by mistake. However, it seems that other proteins with N-terminal extensions blocking the signal peptides are degraded in the cytoplasm.

To confirm the results, three other data sets with peptide evidence for translated upstream translations were analysed. Zhu *et al*. [2] found evidence for 51 N-terminal extensions, and none of the 51 translations were from genes that had signal peptides in their principal isoforms [Figure 2]. Peptides from PeptideAtlas [23] support 63 N-terminal extensions [6]. Here, just one of these extensions preceded a signal peptide [Figure 2]. Finally, Fedorova *et al*. [3] verified 102 N-terminal extensions with an in-house proteomics analysis. Three of the 102 N-terminal extensions (2.9%) were upstream of signal peptides in principal isoforms [Figure 2]. Chi-squared tests for all three analyses found that the number of N-terminal extensions blocking signal peptides that were detected with peptide evidence was significantly lower than would be expected by chance (p < 0.005), confirming that the translation of N-terminal extensions upstream of signal peptides is regulated by the cell.

The Fedorova *et al*. analysis is instructive because the 5’ UTR extensions that it detected can be separated into two groups, those that have accompanying peptide evidence, and those that have just transcript evidence from ribosome profiling experiments. This second group is larger (338 genes) and remarkably 13.6% of the principal isoforms in these genes have signal peptides, very close to the background level in the genome. Together these results confirm that the regulation of proteins that are likely to build up in the wrong cellular compartment takes place during or shortly after translation, suggesting that the *BAG6*-dependent polyubiquitination pathway [11] is the most likely means of degradation.

Other data sets support the Fedorova *et al*. data. Chen *et al*. [24] also detected ribosome profiling evidence (but not peptide evidence) for in-frame upstream translations, this time in 1143 genes. A total of 13.7% of the principal isoforms in these genes were predicted to have signal peptides, close to the percentage for the reference gene set with peptide evidence, and confirming the results from the Fedorova set.

Finally, 262 GENCODE v36 protein isoforms already annotated with extended N-terminals relative to their principal isoforms [4] were analysed. These annotated N-terminal extensions are similar to those detected in the proteomics and ribosome profiling experiments in that the extensions do not have equivalent regions beyond primates, but different in that all the start codons are ATG. A total of 36 genes of these had signal peptides in their principal isoforms, a similar proportion (13.7%) to the background genes detected in proteomics experiments and to the 5’ extensions detected in ribosome profiling experiments.

### Pathogenic variants in N-terminal extensions

The American College of Medical Genetics and Genomics (ACMG) recommends that researchers select the longest or most clinically significant transcript for a gene as the reference transcript [31]. Since there is considerable evidence for transcripts with upstream start codons [3,4,24], many of these regions inevitably will be annotated as coding in gene reference sets and these novel transcripts will almost always become the longest in each gene. So, over time, many researchers will assume that these extended transcripts should be the reference transcript and as a result many will be annotated with pathogenic variants.

This effect can already be seen in some genes. At least two of the genes from the set of 236 annotated non-conserved N-terminal extensions upstream of target peptides in GENCODE v36 are already predicted to have likely pathogenic mutations in the 5’ UTR that produces these extensions. These genes are *TMCO1* [Figure 3], which produces an endoplasmic reticulum-based multipass membrane protein that binds the Sec61 complex [32], and *EDNRB* [Figure 3], a multi-pass transmembrane receptor [33].

**Figure 3.**
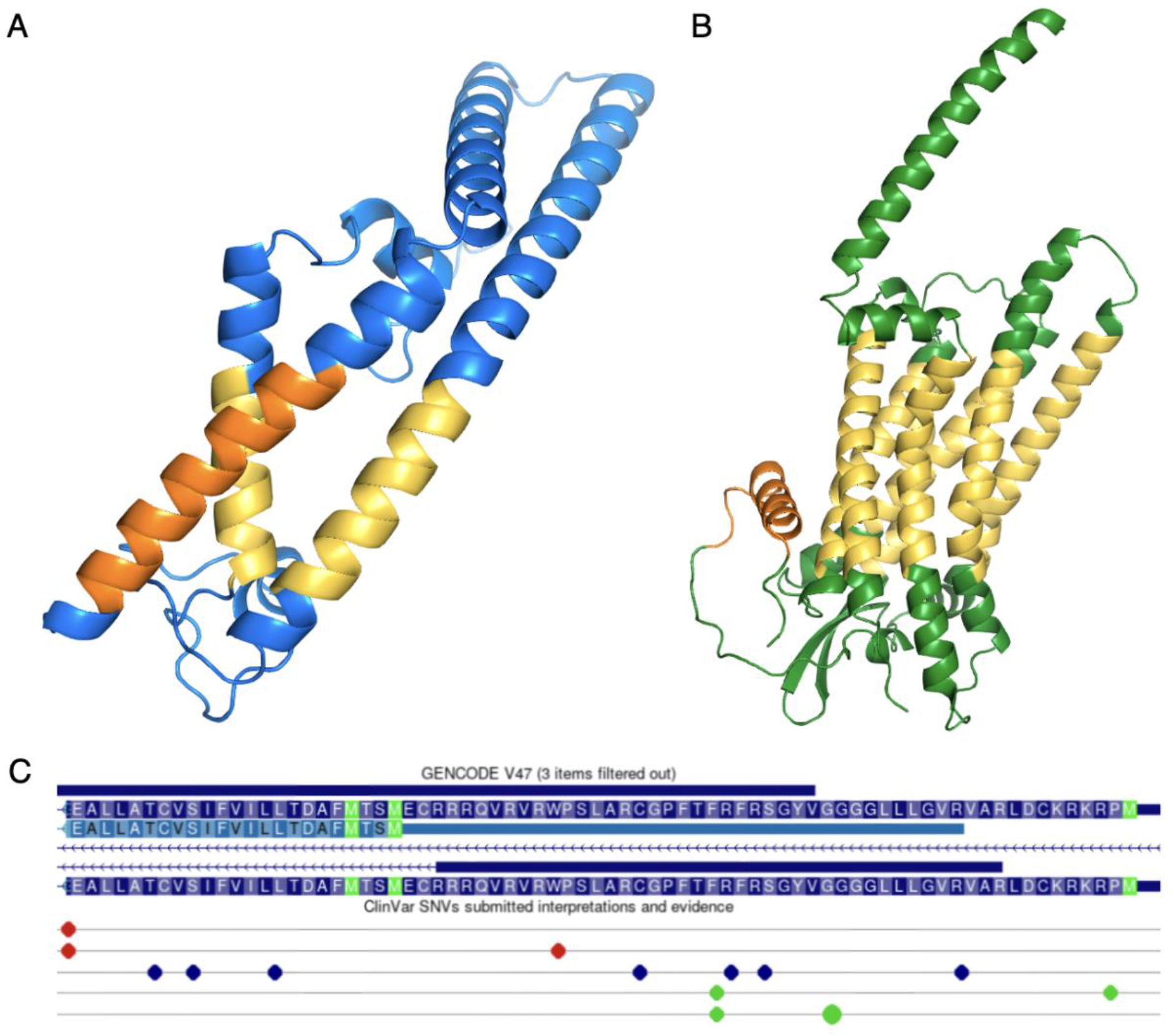
Annotating translated upstream regions attracts pathogenic variants. The likely pathogenic variants annotated in *TMCO1* and *EDNRB*. A. The 3D structure of the principal isoform of *TMCO1* predicted by AlphaFold [29]. Hydrophobic trans-membrane regions coloured in yellow, the hydrophobic region of the predicted signal anchor (uncleaved signal peptide) in orange. B. The AlphaFold 3D structure of *EDNRB*. Hydrophobic trans-membrane regions coloured in yellow, the hydrophobic region of the predicted signal peptide in orange. C. A screenshot of the 5’ exon of *TMCO1* in the UCSC genome browser [34] showing the amino acid sequence, methionines (from ATGs) in green. The methionine on the far right indicates the position of the upstream start codon and the central methionine the canonical start codon. Pathogenic and likely pathogenic variants are shown as red dots below the sequence. The likely pathogenic frame shift variant that affects the upstream region of *TMCO1* is in the centre of the image.

Variants in *TMCO1* can cause craniofacial and skeletal anomalies and glaucoma [35]. The upstream start codon that initiates the translation of the *TMCO1* N-terminal extension is only present in Homininae species and there is no RNASeq evidence for this region in GTex [36], nor peptide evidence in PeptideAtlas. Yet Ensembl/GENCODE and UniProtKB annotate the 5’ extension [Figure 3] and UnIProtKB has even made the extended isoform the representative sequence for this gene. The likely pathogenic variation in the 5’ extension [Figure 3] would lead to a frameshift and is supported by one star in ClinVar [37], but the annotating group has not provided any evidence in its support.

Variants in *EDNRB* may cause Waardenburg syndrome and Hirschsprung disease [38]. The upstream region that produces the N-terminal extension in *EDNRB* is also only intact within Homininae species. There is residual RNASeq expression of this region in GTex, but no reliably identified peptides in PeptideAtlas. Again the extended transcript/isoform is annotated in both Ensembl/GENCODE and UniProtKB and the likely pathogenic variation in the 5’ extension (a premature stop codon) has one star in ClinVar.

## Conclusions

Evidence that 5’ UTRs upstream of canonical start codons are being translated is indisputable [3,24] and an ever increasing number of studies find peptide support for these upstream translations [1-4,6]. The most common form of upstream translation is the in-frame translation of 5’ UTR that produces N-terminally extended canonical protein isoforms [4,6].

This analysis finds clear evidence that at least some of these N-terminally extended protein isoforms are under regulatory control and are degraded by the cellular machinery to prevent the build up of secretory and trans-membrane proteins in the cytoplasm where they might be harmful to the cell.

Protein isoforms with extended N-terminals have additional amino acids at the N-terminal end, and if the protein has a signal peptide, the extra amino acids would have the effect of moving it further away from the N-terminal. This would mean it would be more likely that the signal peptide would still be in the ribosome tunnel when the signal recognition particle disengages from the ribosome [10]. Once the signal recognition particle disengages from the ribosome, the protein will be destined for the cytoplasm rather than the endoplasmic reticulum [11,14]. Unless it is recognised by cellular surveillance and tagged for degradation, the N-terminally extended isoform will accumulate in the cytoplasm.

In each of the four unrelated proteomics analyses that were analysed, N-terminal extensions that would abolish signal peptides were detected significantly less frequently than would be expected, suggesting that these N-terminally extended isoforms are degraded. Since the translated signal peptide would leave an exposed hydrophobic region, the most likely path to degradation would involve *BAG6*, which has been shown to promote the degradation of proteins with N- and C-terminals that have substantial hydrophobic regions [11,14].

Degradation of the mislocalised proteins was confirmed to occur post-translation because the 5’ extensions that were detected in ribosome profiling experiments [3, 24] had the same proportion of signal peptides as all genes in the control set, those genes detected in proteomics experiments [4]. When we compared compendiums of the four large-scale proteomics analyses and the two ribosome profiling experiments, there was more than eight times as much support for upstream regions that would block signal peptides among the ribosome profile transcripts.

Given that translation from upstream start codons is so frequent [3,4,6,24], one important question is whether or not there are any adaptive benefits. Whether these upstream translations have gained functional roles [39], or whether the translations are simply the result of noisy translation initiation and are not under selection pressures [40]. The simple answer seems to be, both but mostly the result of inefficient translation initiation.

Novel translated upstream regions have a series of characteristics that are not indicative of coding regions [4]. Most importantly, a large majority of upstream translations are not conserved among primates [3,4]. There is no evidence of purifying selection in germline variation data [41] from the majority of upstream regions that are not conserved across primates [4]. At the same time, a small proportion, including the members of the C1QL family, has strong cross species conservation and does appear to be under selection pressure [4].

The results from this analysis demonstrate that practically all the N-terminal extensions that precede signal peptides are degraded. If they are degraded, it would seem to confirm that this sub-group of non-conserved upstream translations are potentially harmful biological aberrations. Since the production of harmful biological aberrations is not the result of any evolutionary selection, it is almost certainly a side-effect of a noisy translation initiation process.

If one sub-group of unconserved N-terminal extensions are a side-effect of a noisy translation initiation process, then this is almost certainly also true for the rest of the non-conserved N-terminal extensions. The only difference being that these N-terminally extended isoforms are not degraded because they are produced in small quantities in the correct cellular compartment. They are more likely to be tolerated by the cell. The degradation of regions translated from upstream start codons confirms that translation initiation is an especially noisy biological process.

The observation that non-canonical ATGs and their associated Kozak sequences are selected against in all frames upstream and downstream of the canonical ATG [42] is another indication that translation initiation is prone to molecular errors. Since ATGs are much more efficient translation initiation codons than non-ATG start codons [43], accidental translation from a non-canonical ATG has the potential to be much more of a metabolic burden [42]. The inefficiency of translation from non-canonical start codons may be one of the reasons that so much upstream translation is tolerated by the cell.

If the majority of non-conserved upstream translations are the result of molecular errors, there has to some consideration as to how they are annotated in reference gene sets to avoid the risk of being annotated with unsupported pathogenic variants like *TMCO1* and *EDNRB* and propagating errors.

Although the ACMG still recommends that researchers select the longest transcript for a gene as the reference transcript [31], the MANE Select [44] transcript is now available for more recent versions of reference gene sets. While this will go some way to avoid the circular annotation of unsupported pathogenic variants, it is not a panacea, partly because protein length and pre-existing pathogenic variantions were important inputs in this automatic process.

One extreme case of what could happen can be seen in the gene *PUS1*. The MANE Select is the longest transcript, generated from an upstream start codon, presumably because an unsupported pathogenic variant (a premature stop) was already annotated in the upstream exon. However, the region between the upstream start codon and the canonical start codon has little transcript support and is not under coding conservation, even in primates. At present, the *PUS1* upstream region is annotated with seven pathogenic or likely pathogenic variants in ClinVar, most of which have been added since it was tagged as part of the MANE Select transcript and none of which have any support, save that they are high impact variants and are present in a gene known to play a role in a rare mitochondrial myopathy [45]. One of the submitters was swayed to record the variant as pathogenic because of the ClinVar pathogenic variants [37] already present in the exon, the very definition of circular annotation.

A definition of functional importance that includes conservation would help distinguish canonical and conserved alternative isoforms from those isoforms that are the result of imprecise translation initiation, but no reference gene set incorporates any such method at present. The APPRIS database includes TRIFID functional scores [46] for each isoform, which give an idea of the relative conservation. High scoring TRIFID isoforms have been shown to capture almost all reliably annotated pathogenic variants in ClinVar [21].

This analysis shows that degradation by the ubiquitination pathway is the likely fate of erroneous upstream translation initiation when it leads to the blocking N-terminal localization signals. That a large proportion of translations from upstream start codons appear to be products of a misfiring translation initiation process has implications for the huge numbers of upstream translations that are being detected in large-scale analyses, and strongly suggests that further work needs to be carried out in this area.

## Acknowledgemnents

This work was funded by the National Human Genome Research Institute of the National Institutes of Health (grant number U41 HG007234). The author would like to thank Federico Abascal for his input on the paper.

